# Histone H4K20 Trimethylation is Decreased in Murine Models of Heart Disease

**DOI:** 10.1101/2022.02.10.479885

**Authors:** Samuel M. Hickenlooper, Kathryn Davis, Hanin Sheikh, Mickey Miller, Ryan Bia, Steven Valdez, Marta W. Szulik, Sarah Franklin

## Abstract

Heart disease is the leading cause of death in the developed world, and its comorbidities such as hypertension, diabetes, and heart failure are accompanied by major transcriptomic changes in the heart. During cardiac dysfunction, which leads to heart failure, there are global epigenetic alterations to chromatin that occur concomitantly with morphological changes in the heart in response to acute and chronic stress. These epigenetic alterations include the reversible methylation of lysine residues on histone proteins. Lysine methylation on histone H3K4 and H3K9 were among the first methylated lysine residues identified and have been linked to gene activation and silencing, respectively. However, much less is known regarding other methylated histone residues, including histone H4K20. Trimethylation of histone H4K20 has been shown to repressive gene expression, however this mark has never been examined in the heart. Here we utilized immunoblotting and mass spectrometry to quantify histone H4K20 trimethylation in three models of cardiac dysfunction. Our results show that lysine methylation at this site is regulated in a biphasic manner leading to increased H4K20 trimethylation during acute hypertrophic stress and decreased H4K20 trimethylation during sustained ischemic injury and cardiac dysfunction. In addition, we examined publicly available datasets to analyze enzymes that regulate H4K20 methylation and identified one demethylase (KDM7C) and two methyltransferases (KMT5A and SMYD5) which were all upregulated in heart failure patients. This is the first study to examine histone H4K20 trimethylation in the heart and to determine how this post-translational modification is differentially regulated in multiple models of cardiac disease.

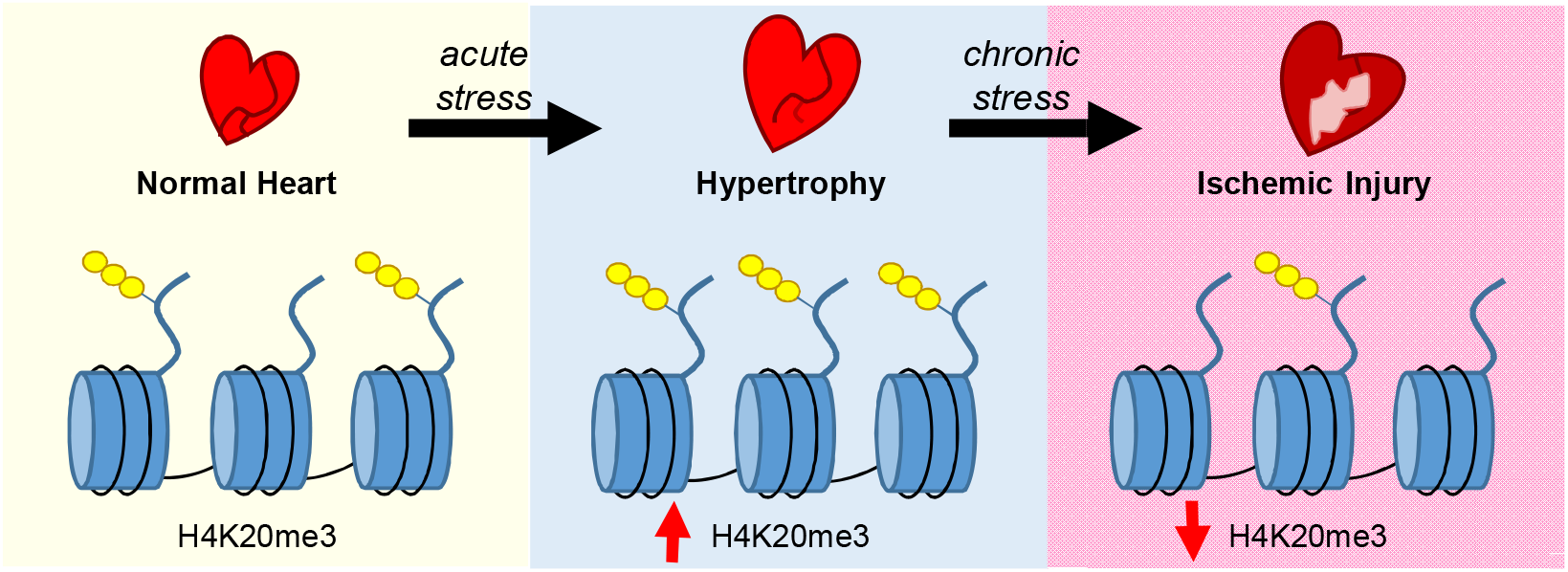

## INTRODUCTION

Heart disease is the leading cause of death throughout the world.(1) Precursors to heart disease include common conditions such as diabetes and hypertension. These persistent stressors can lead to large changes in cardiac morphology, and often cause many types of cardiac dysfunction including heart failure. It has also been shown that many cardiac pathologies result from dynamic changes in gene expression and conversely that modulating epigenetic factors in murine models can prevent or abrogate ischemic injury and pathological remodeling.(2-4) For example, during cardiac hypertrophy and failure, there is global upregulation of genes more commonly involved in heart development.(5, 6) These largescale transcriptome changes are preceded by epigenetic alterations to chromatin.

Chromatin contains DNA wrapped around an octamer of histone proteins (two copies each of H2A, H2B, H3, and H4) which have N-terminal tails that extend beyond the core protein structure and interact with a large number of regulatory proteins.(7) Transcription can be activated or silenced depending on the chromatin environment: for example, more accessible euchromatin, allows transcription factors and other transcriptional machinery to bind chromatin, while less accessible heterochromatin, prevents this binding and silences gene expression.(8) One epigenetic mechanism which contributes to the chromatin landscape and gene expression is the post-translational modification (PTM) of histone proteins.(9, 10) Histones are subject to a wide number of post-translational modifications including the methylation of arginine(R) and lysine(K) residues. Regarding lysine methylation, this amino acid can be reversibly methylated and can accept up to three methyl groups (mono-, di-, or tri-methylation, respectively, abbreviated as me1, me2 and me3).(9, 11) Sites of lysine methylation, including K4 and K9 on histone H3, were among the first methylated lysine residues to be identified(12-14) and have been linked to gene activation and silencing, respectively. While these two residues have been a main focus of research regarding lysine methylation and its role in transcriptional regulation, much less is known regarding other methylated lysine residues. Among the less studied modifications is histone H4K20 methylation.

The methylation of histone H4K20 was first discovered by DeLange et al. in which they examined residues of histone tails in calf and pea extracts, however the exact function of this histone mark was not known at the time.(15) Three decades later, the first reports were published that linked specific methyltransferases responsible for methylation of histone H4 at lysine K20 together with functional studies that revealed the effects of histone H4K20 methylation on chromatin status and transcriptional regulation of genes.(16-19) In general, methylation of H4K20 has been implicated in gene silencing, heterochromatin formation, mitosis and genomic stability.(20-22) More specifically, mono- and di-methylation of H4K20 are transcriptional activating marks implicated in cell cycle regulation and DNA repair(23, 24) while trimethylation is most often associated with heterochromatin formation and gene repression.(20) Most commonly, H4K20me3 is located at gene promoters and functions to repress transcription alongside specific chromatin readers. Each of these methylation states are carried out by separate enzymes called lysine methyltransferases and these methyl marks are selectively removed by lysine demethylases (Table 1).(25-27) Notably, there are only six enzymes that catalyze or remove H4K20 trimethylation that have been identified to date (visualized in Figure 1) compared to sixteen total enzymes that regulate H3K4 trimethylation.(28) This suggests that there may be other enzymes that regulate H4K20 trimethylation that have not been identified to date. While we are beginning to understand the complexity of these methylation marks and the diverse processes they regulate, much less is known regarding their role in specific cell types and pathologies. Along these lines, the methylation of histone H4K20 has never been examined in the heart or cardiomyocytes.

**Table 1.**
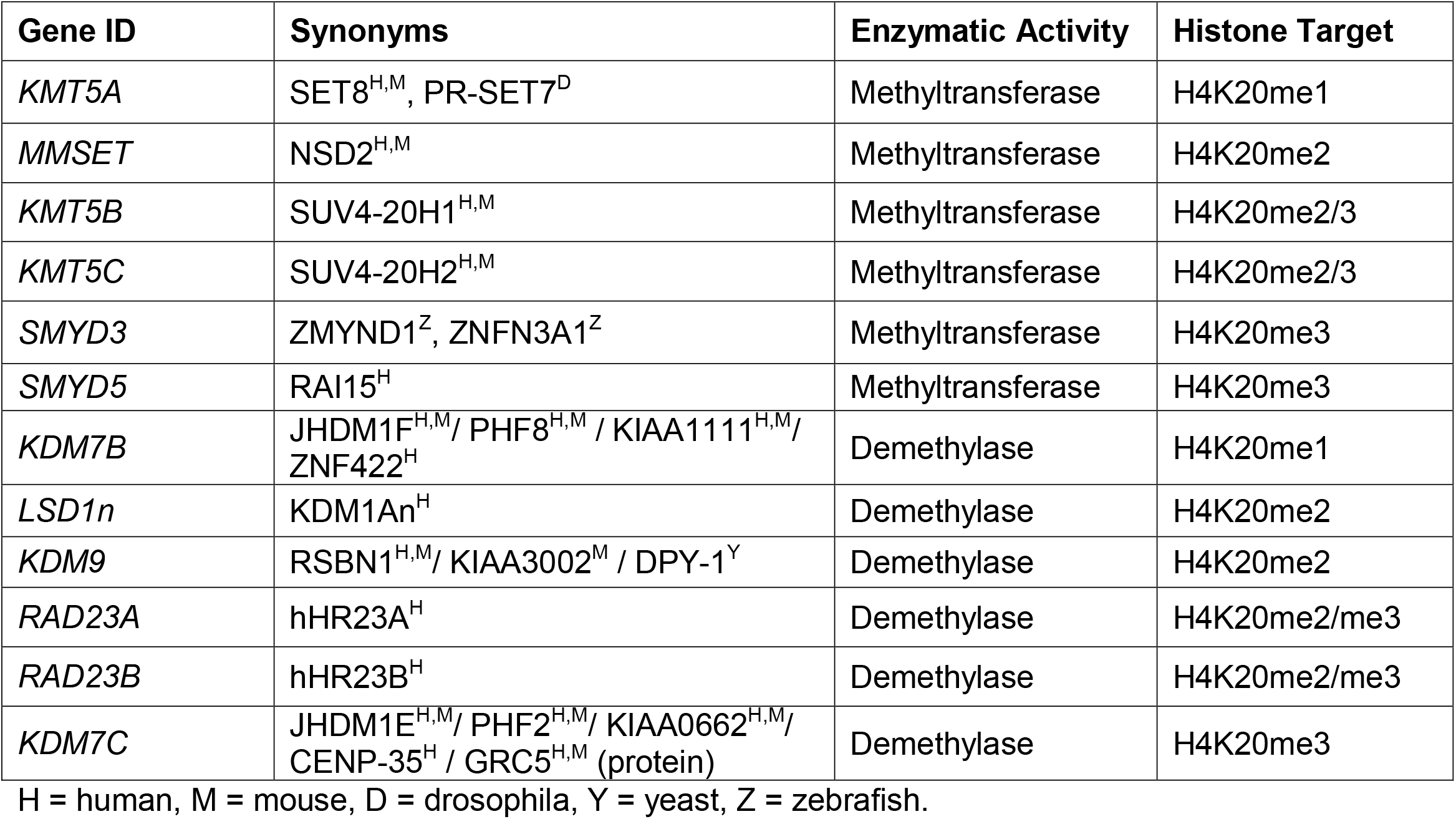
Enzymes That Regulate H4K20 Methylation

| Gene ID | Synonyms | Enzymatic Activity | Histone Target |
| --- | --- | --- | --- |
| *KMT5A* | SET8<sup>H,M</sup>, PR-SET7<sup>D</sup> | Methyltransferase | H4K20me1 |
| *MMSET* | NSD2<sup>H,M</sup> | Methyltransferase | H4K20me2 |
| *KMT5B* | SUV4-20H1<sup>H,M</sup> | Methyltransferase | H4K20me2/3 |
| *KMT5C* | SUV4-20H2<sup>H,M</sup> | Methyltransferase | H4K20me2/3 |
| *SMYD3* | ZMYND1<sup>Z</sup>, ZNFN3A1<sup>Z</sup> | Methyltransferase | H4K20me3 |
| *SMYD5* | RAI15<sup>H</sup> | Methyltransferase | H4K20me3 |
| *KDM7B* | JHDM1F<sup>H,M</sup> / PHF8<sup>H,M</sup> / KIAA1111<sup>H,M</sup> / ZNF422<sup>H</sup> | Demethylase | H4K20me1 |
| *LSD1n* | KDM1An<sup>H</sup> | Demethylase | H4K20me2 |
| *KDM9* | RSBN1<sup>H,M</sup> / KIAA3002<sup>M</sup> / DPY-1<sup>Y</sup> | Demethylase | H4K20me2 |
| *RAD23A* | hHR23A<sup>H</sup> | Demethylase | H4K20me2/me3 |
| *RAD23B* | hHR23B<sup>H</sup> | Demethylase | H4K20me2/me3 |
| *KDM7C* | JHDM1E<sup>H,M</sup> / PHF2<sup>H,M</sup> / KIAA0662<sup>H,M</sup> / CENP-35<sup>H</sup> / GRC5<sup>H,M</sup> (protein) | Demethylase | H4K20me3 |

H = human, M = mouse, D = drosophila, Y = yeast, Z = zebrafish.

**Figure 1.**
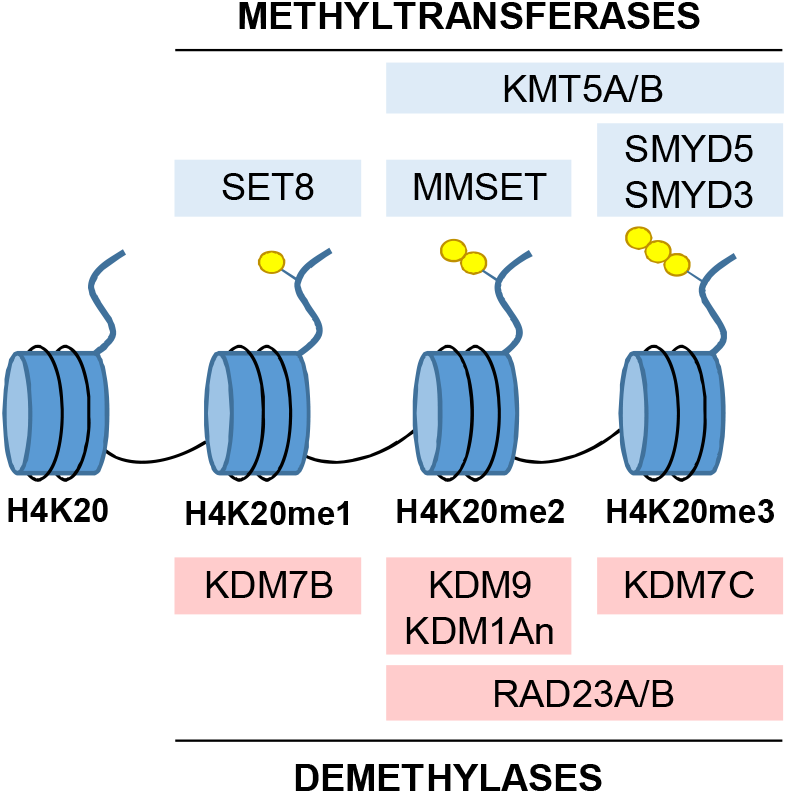
Enzymes that regulate H4K20 methylation. There are relatively few enzymes known to regulate H4K20 methylation. Blue boxes indicate methyltransferases that catalyze each methylation state. Red boxes indicate demethylases that remove specific methyl marks from H4K20 in sequential order. (Yellow circles depict H4K20 methylation).

Analysis of global changes in histone post-translational modifications (versus methylation on specific genes) can provide insights into how the transcriptome changes in response to stress or disease. Presently, there have only been three studies which have examined the total abundance of histone methylation in the context of heart disease. These studies examined histone H3K4me3(29, 30), H3K9me3(29, 30), and H3K36me3(31) in human heart failure patients and rodent models of heart failure and showed global increases in H3K4me3 abundance and decreases in H3K9me3 in cardiac tissue.(29, 30) In addition, studies in both rats and humans, using ChIP-Seq-based profiling showed that the genomic regions enriched in these marks changes substantially in failing hearts compared to healthy donor controls, indicating a significant shift in which genes were being actively transcribed.(29, 31) These results suggest that modulation of histone methylation is important in the development and progression of heart disease. However, studies regarding other sites of methylation on histones, including H4K20me3, have not been examined in the healthy or diseased heart.

In this study, we utilized both cell and animal models of cardiac dysfunction to examine histone H4K20 trimethylation abundance during disease. Our results show H4K20 trimethylation is increased in cardiomyocytes during acute hypertrophic stress but is decreased in the heart after long-term exposure to isoproterenol or ischemic injury in mice. In addition, we profiled public databases of methyltransferases and demethylases which regulate histone H4K20 methylation to determine which of these enzymes are differentially expressed in human heart failure patients. Specifically, we identified one demethylase (KDM7C) and two methyltransferases (KMT5A and SMYD5) which were all upregulated in heart failure patients. This is the first study to examine histone H4K20 trimethylation in the heart and to determine how this post-translational modification is differentially regulated in multiple models of cardiac disease.

## METHODS

### Mouse Models of Cardiac Stress

All protocols involving animals conform to the NIH Guide for the Care and Use of Laboratory Animals and were approved by the Institutional Animal Care and Use Committee of the University of Utah. All efforts were made to minimize pain and distress during procedures and isolation of the heart by anesthesia. In this study, we used both male and female FVB mice.

### Isoproterenol Treatment of Mice

FVB mice between the age of 8-12 weeks were treated with isoproterenol (15 mg/kg/d) via subcutaneous implantation of osmotic mini-pumps (Alzet 2004) for 6 weeks as previously described.(32, 33) Briefly, pumps were prepared before implantation by filling the reservoir with sterile isoproterenol solution (saline was used as a control). Mice were anesthetized, and subcutaneous tissue on the back was spread apart by inserting a hemostat to create enough room for the pump. The filled pump was inserted and kept clear of vital organs and the incision. The incision was closed with a sterile suture. Cardiac tissue was harvested at 6 weeks, weighed and used for downstream analyses.

### Ischemic Injury via Left Anterior Descending Coronary Artery Ligation

The left anterior descending coronary artery (LAD) was ligated in FVB mice as reported previously with minor modifications.(34) Briefly, mice were put under mechanical ventilation with anesthesia and the chest was opened through a left thoracotomy. A suture was placed around the left anterior descending coronary artery, as it emerges from under the left atrium, to permanently occlude the LAD. The sham control group underwent the same surgical procedure without ligation of the LAD. Mice were sacrificed three weeks after LAD ligation and cardiac tissue was excised, weighed and used for downstream analyses.

### Cultured H9c2 Cells (Rat Cardiomyoblasts)

H9c2 cells were cultured in Dulbecco’s Modified Eagle Medium (DMEM) supplemented with 10% Fetal Bovine Serum (FBS) and 1% penicillin-streptomycin (P/S) until 80% confluent. Cells were then differentiated in DMEM supplemented with 1% FBS and 1% P/S for 48 hours. To induce hypertrophic growth cells were treated with 100 µM phenylephrine (PE) for 48 hours and then harvested for experiments.

### Cultured 3T3 Mouse Fibroblasts

3T3 fibroblasts were cultured in Dulbecco’s Modified Eagle Medium (DMEM) supplemented with 10% Fetal Bovine Serum (FBS) and 1% penicillin-streptomycin (P/S) until 80% confluent. To compare histone methylation in these cells to cardiomyocyte cells in hypertrophic stress they were similarly treated with 100 µM phenylephrine (PE) for 48 hours and then harvested for experiments.

### Western Blotting

Protein samples were diluted in Laemmli buffer, resolved on 12% SDS-PAGE gels, and then transferred to nitrocellulose membranes using the semi-dry transfer method (Bio-Rad) as we have previously published.(35, 36) The success of protein transfer to the membrane was confirmed using Ponceau S (0.1% w/v in 5% acetic acid). These membranes were blocked with either 5% milk in TBS-T or 5% BSA in TBS-T, depending on the suggestions from the manufacturer (Abcam). Antibodies used in this study are as follows: Histone H4K20me3 (Abcam, ab9053), Histone H4 (Abcam, ab10158), β-Tubulin loading control (Abcam, ab6046), and goat anti-rabbit (Abcam, ab97051).

### Quantitative Real Time PCR Analysis

Total RNA was isolated with Trizol according to the manufacturer’s instructions, as we have previously published.(37) cDNA was synthesized using the Superscript III First Strand Synthesis System (Life Technologies) and PCR was performed using a Bio-Rad CFX Connect RT-PCR detection system. The primers used were for *Nppa* (FWD: CTGATGGATTTCAAGAACCTGCT, REV: CTCTGGGCTCCAATCCTGTC) as a marker of cardiac stress and myocyte hypertrophy and *β-Actin* (FWD: GGGGTGTTGAAGGTCTCA, REV: TGTTACCAACTGGGACGA) as a housekeeping gene.

### Liquid Chromatography-Tandem Mass Spectrometry

Sample preparation for mass spectrometry was performed using a standard FASP protocol that we have previously published.(30) Samples were diluted with Urea buffer, reduced, and alkylated, then proteins were digested with trypsin overnight at 37°C, and acidified using 1% formic acid.

Peptides were analyzed with an Orbitrap Velos Pro mass spectrometer (Thermo) interfaced with an Easy nLC-1000 UPLC and outfitted with a ESI Source Solutions pulled tip column type 2 (15 cm x 75 μm inner diameter, 3 μm particle size, 120 Å pore diameter, ESI Source Solutions). Spectra were acquired in data-dependent mode with dynamic exclusion enabled and peptides were fragmented using CID fragmentation. The top twenty MS1 peaks were analyzed at a resolution of 30,000. Samples were run in duplicate to generate technical replicates.

The resulting spectra were analyzed using MaxQuant 1.6.7.0 against the UniprotKB mouse protein database. Search engine parameters for MaxQuant were as follows: trypsin digestion, two missed cleavages, precursor mass tolerance of 20 ppm, and fragment mass tolerance of 0.5 Da. Match between runs was enabled with a match time window of 0.3 minutes, match ion mobility window of 0.05 and an alignment time window of 20 minutes. The false discovery rate (FDR) was 1%. For relative quantification, peptide abundance (measured as intensity values for area under the curve) was used to calculate the abundance or total histone H4 (7 mapped peptides) and Gapdh (15 mapped peptides). Normalized log2 intensities of peptides were used in statistical comparisons of groups: Student’s two sample t-test was used for comparisons between the two sample groups and statistical analyses were performed in Perseus 1.6.5.0. The mass spectrometry proteomics data have been deposited to the PRIDE Archive (http://www.ebi.ac.uk/pride/archive/) via the PRIDE partner repository.

## RESULTS

### Isoproterenol treatment in mouse hearts leads to decreases in H4K20 trimethylation

Cardiac dysfunction is preceded by large transcriptomic changes within the heart, which presumably involves changes in histone post-translational modifications. To examine whether H4K20 trimethylation levels change in the diseased heart, we first examined cardiac tissue from mice after isoproterenol treatment, which is a β-adrenergic receptor agonist that induces cardiac hypertrophy and failure.(38) This model mimics non-ischemic cardiomyopathy, as seen by chronic adrenergic stimulation due to elevated catecholamine levels, in heart failure patients.(33, 39, 40) We implanted mice with isoproterenol-filled osmotic mini-pumps that allows continuous diffusion of isoproterenol, or saline as a control, over the course of 6 weeks (Figure 2A). At this time point mice were experiencing cardiac hypertrophy, confirmed by increased expression of *Nppa* in the heart via qPCR (Figure 2B). We also quantified changes in histone H4K20 trimethylation in cardiac tissue and observed a ∼30% decrease in abundance (Figure 2C, D).

**Figure 2.**
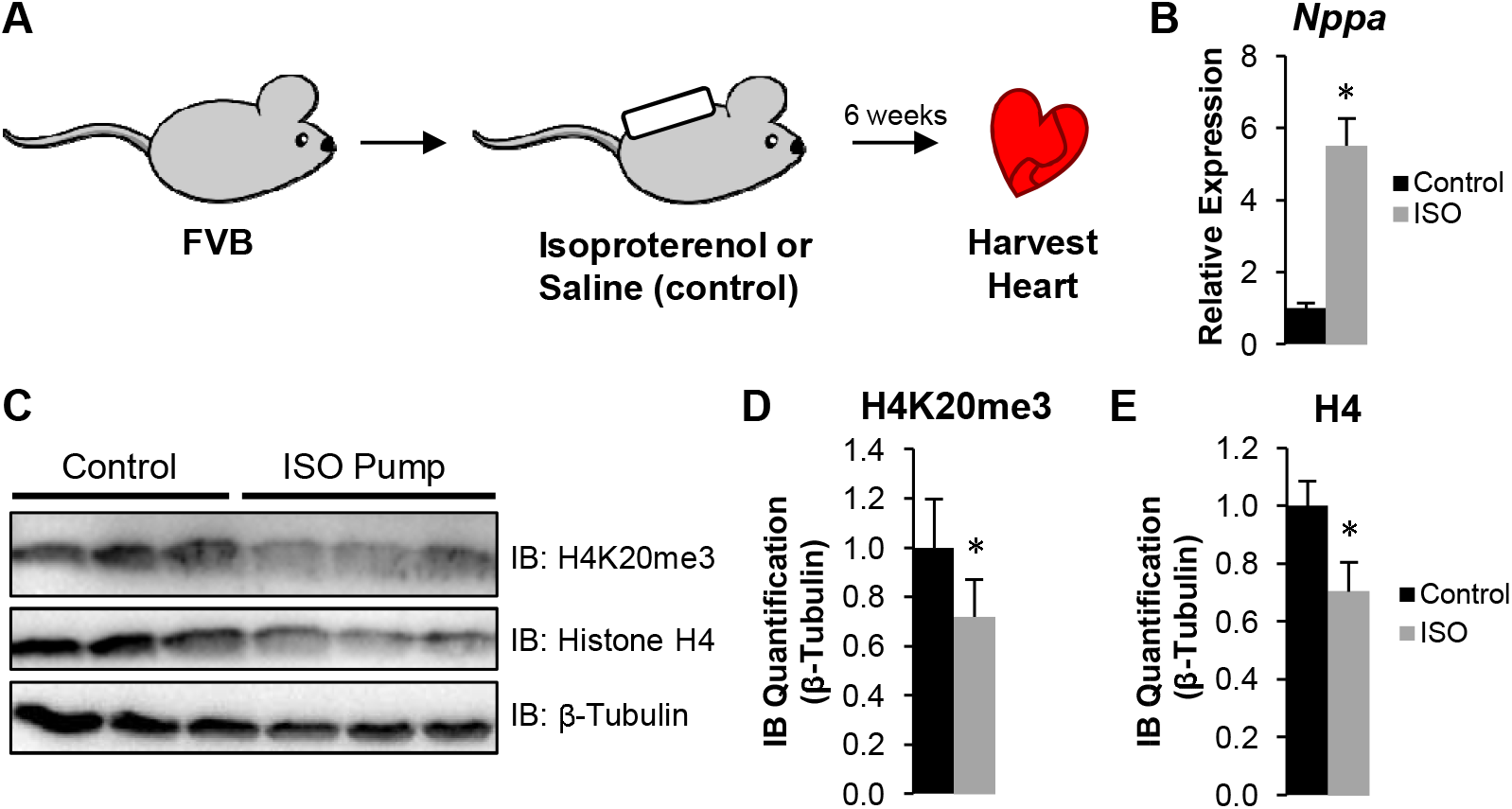
Decreased H4K20 trimethylation and histone H4 in mouse heart after isoproterenol-induced cardiomyopathy. **A)** Mice were treated with isoproterenol via mini-osmotic pump implantation or saline (control) for 6 weeks. (n=3) **B)** Cardiac stress was confirmed by quantifying Nppa via RT-qPCR. **C)** Immunoblot analysis of cardiac tissue from these animals showed a decrease in **(D)** histone H4K20 trimethylation and **(E)** histone H4 (both normalized to β–Tubulin). Asterisk indicates p-value <0.05.

### Ischemic injury via LAD ligation in mice results in decreased H4K20 trimethylation

To evaluate levels of histone H4K20me3 in a separate mouse model, we performed permanent ligation of the LAD in the heart (Figure 3A), which mimics ischemic cardiomyopathy via myocardial infarction.(41) To confirm cardiac injury, we measured *Nppa* in cardiac tissue via qPCR which increased 15-fold (Figure 3B). Then we examined the abundance of histone H4K20me3 and total histone H4 via western blotting which decreased ∼70% and 30%, respectively, when compared to β-Tubulin (loading control) (Figure 3C-F). Because of two key reasons, we decided to quantify histone H4 abundance using an additional technique (via mass spectrometry): 1) post-translational modifications have been reported to effect the efficiency of antibody binding in antibody-based assay,(42, 43) and 2) previous reports have suggested that global histone H4 levels are regulated at both the transcriptional and post-transcriptional levels to keep them uniquely stable and tightly controlled(44). Quantification of histone H4 by mass spectrometry (using 7 unique peptides) and normalized to Gapdh (using 15 unique peptides) showed no significant change in abundance (Figure 3G).

**Figure 3.**
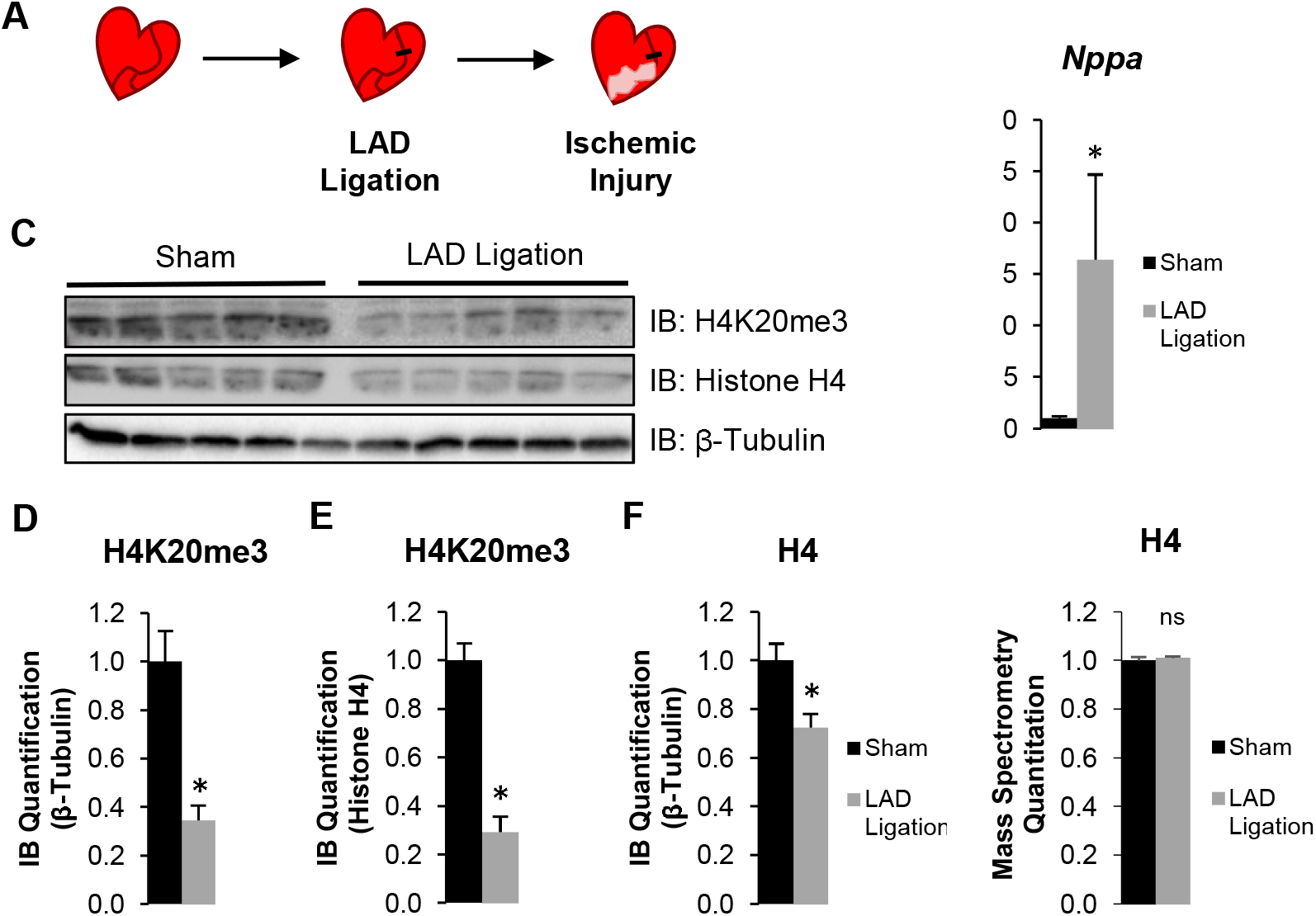
Decreased H4K20 trimethylation in mouse heart after ischemic injury. **A)** Mice were subjected to ischemic injury by ligating the left anterior descending (LAD) coronary artery. Sham surgery was used as a control where no ligation was performed. (n=5) **B)** Cardiac tissue was excised and Nppa was quantified via RT-qPCR to confirm cardiac dysfunction. **C)** Immunoblotting of cardiac tissue showed a decrease in H4K20 trimethylation when normalized to **(D)** β-Tubulin or **(E)** histone H4 abundance. Global abundance of histone H4 was decreased in cardiac tissue when examined by **(F)** immunoblotting but not by **(G)** mass spectrometry. Asterisk indicates p-value <0.05. (ns= not significant)

### Phenylephrine-induced hypertrophy in isolated cardiomyocytes increases H4K20 trimethylation

To investigate whether histone H4K20 trimethylation changes in a cell model of hypertrophy we cultured H9c2 cells (cardiomyoblasts), differentiated them in low-serum media to confer a more cardiomyocyte-like phenotype, and treated them with phenylephrine (PE), an α-adrenergic receptor agonist (Figure 4A). After 48h we observed increased expression of *Nppa* (∼5-fold), an established marker of cardiac hypertrophy (Figure 4B). Quantification of histone H4K20 trimethylation in these cells (Figure 4C-E) showed a 1.8-fold increase with no significant change in histone H4 abundance. We also examined changes in H4K20 trimethylation in fibroblasts after PE treatment, as they are also abundant in the heart. Our results show a ∼3-fold increase in H4K20 trimethylation with no significant change in histone H4 abundance (Figure 4F-H).

**Figure 4.**
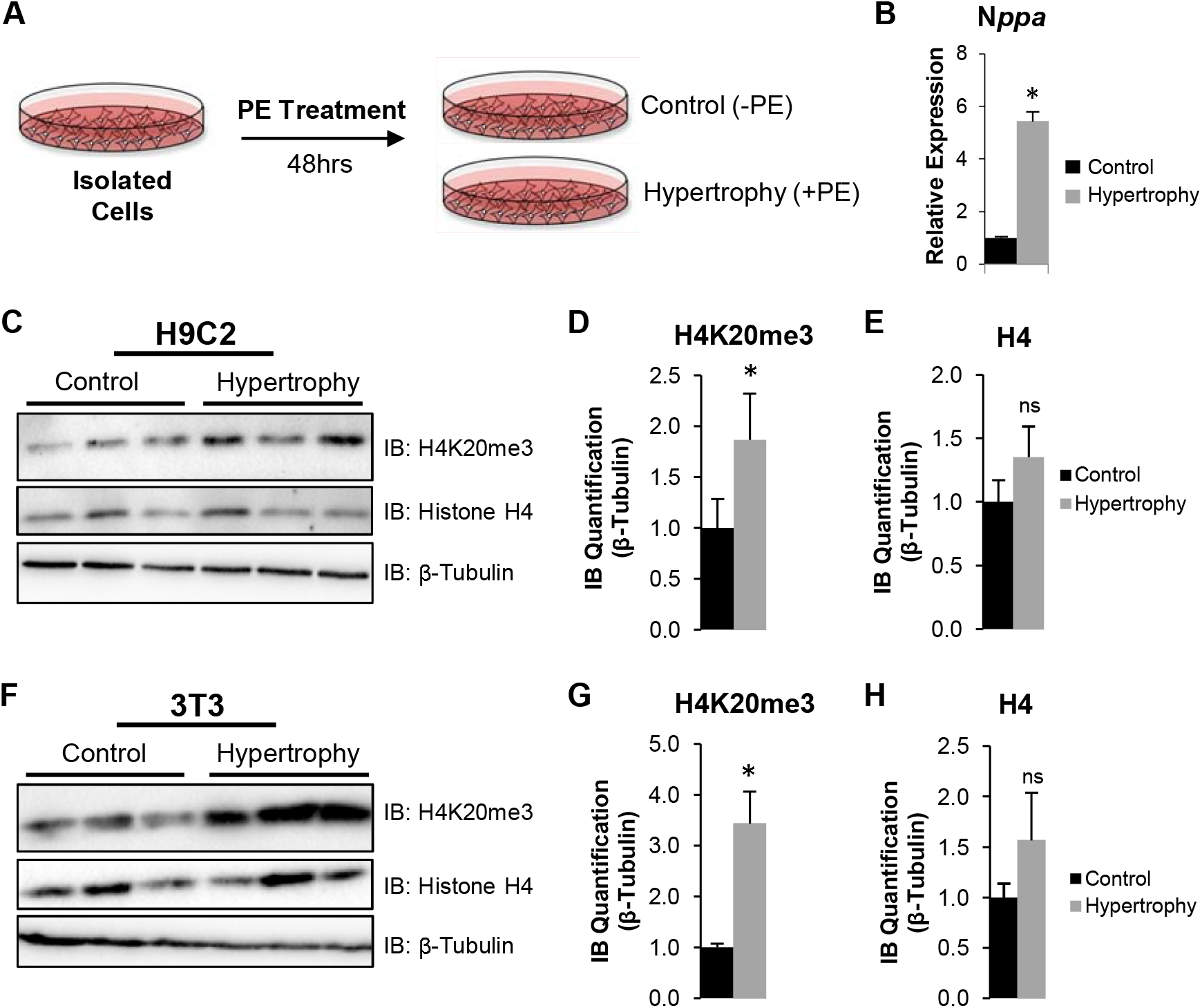
Acute hypertrophy in isolated cells increases H4K20me3 abundance. **A)** Cultured cardiomyocytes (H9c2 cells) and 3T3 fibroblasts were treated with phenylephrine (PE) for 48 hrs. (n=3) **B)** Relative expression of Nppa in H9c2 cells via RT-qPCR confirmed hypertrophic treatment. **C)** Immunoblotting of H9c2 cell lysates showed an increase in **(D)** histone H4K20 trimethylation with no change in **(E)** total histone H4 abundance (both normalized to β-tubulin). **F)** Immunoblotting of 3T3 cell lysates showed an increase in **(G)** histone H4K20 trimethylation with no change in **(H)** total histone H4 abundance (both normalized to β-tubulin). Asterisk indicates p-value <0.06. (ns= not significant)

### Enzymes that regulate H4K20 methylation are differentially expressed in human heart failure

The changes we observed in histone H4K20 methylation in cardiac tissue must be accomplished by the enzymes which regulate these methyl marks, primarily lysine methyltransferases and demethylases. Therefore, we examined publicly available datasets from three separate studies which analyzed transcript expression in cardiac tissue from patients experiencing ischemic cardiomyopathy (ICM), non-ischemic cardiomyopathy (NICM), or dilated cardiomyopathy (DCM), Sweet et al.(45), Yang et al.(46), Liu et al.(47). For each study statistical significance was determined on an individual study basis using the filters published which included a fold change equal to or greater than 1.2 with a p-value less than 0.05. In these studies, we found three enzymes which regulate H4K20 methylation were differentially expressed in at least one disease condition (Table 2). Specifically, one demethylase, KDM7C, was upregulated 1.43-fold in ischemic cardiomyopathy and 2.2-fold in dilated cardiomyopathy, and two methyltransferases (KMT5A and SMYD5) were upregulated 1.2-fold in ischemic cardiomyopathy and non-ischemic cardiomyopathy, respectively.

**Table 2.** H4K20 Methyltransferases and Demethylases Differentially Expressed in Human Cardiac Disease

| Gene ID | All Known Histone Targets | ICM | NICM | DCM | Cardiac KO/KD Phenotype |
| --- | --- | --- | --- | --- | --- |
| *KMT5A/ SET8/ PR-SET7* | H4K20me1 | +1.2(47) |  |  | Unknown |
| *KDM7C/ PHF2/ JHDM1E* | H3K9me1/me2, H4K20me3 | +1.43(45) |  | +2.2(47) | Unknown |
| *SMYD5* | H4K20me3 |  | +1.2(46) |  | *Zebrafish*: Normal cardiac development(50) |

Enzymes that are differentially expressed in human cardiac disease. Values indicate fold change for transcripts with a $P \leq 0.05$ . Negative value corresponds to downregulated transcripts and positive value to upregulated transcripts. ICM, ischemic cardiomyopathy; NICM, nonischemic cardiomyopathy; DCM, dilated cardiomyopathy.

## DISCUSSION

Here, we provide the first evidence of differential regulation of histone H4K20 trimethylation in multiple models of cardiac dysfunction (summarized in the graphical abstract). Our results show an increase in H4K20 trimethylation in cultured cells subjected to acute hypertrophic stress while mouse models of sustained cardiac dysfunction show a marked decrease. This biphasic regulation suggests different stress response mechanisms during acute and chronic stress and are consistent with other signaling pathways in these models. This post-translational modification has been primarily linked to gene repression (in non-cardiac cells) and suggests that this modification, and the associated enzymes, are key regulators of gene expression in cardiomyocytes during stress. These results, showing a global decrease in this repressive histone mark, are consistent with previous findings showing a global decrease in H3K9me3 (mark of gene repression) and increase in histone H3K4me3 (mark of gene activation).(48) Indeed, it has been suggested that global changes in gene expression patterns in cardiomyocytes may favor a more undifferentiated or fetal-like gene expression pattern,(49) often characterized by euchromatin, and enable the cell to dynamically adapt to stress. While this is the first study to correlate global changes in H4K20me3 abundance with the development of heart disease, additional analyses will be needed to identify the specific genes regulated by this post-translational modification in the cardiomyocyte and their involvement in cardiac pathologies.

In addition, we identified two methyltransferases (KMT5A, SMYD5) and one demethylase (KDM7C) whose expression is altered in human heart failure patients. Of these three enzymes, only one has been examined in the heart previously, in a single study which showed that loss of Smyd5 in the developing zebrafish had no effect on cardiac development.(50) No studies have ever examined these three enzymes in the adult heart.

One additional observation in this data is the decrease in total histone H4 levels seen by immunoblotting of cardiac tissue in both mouse models (albeit to varying degrees). Very few studies have reported dynamic changes in total histone protein levels which have previously only been seen in non-cardiac cells.(51) This discrepancy between histone H4 levels observed by immunoblotting versus mass spectrometry may be due to impaired antibody binding caused by unknown neighboring histone post-translational modifications. This phenomenon has been previously reported as a major challenge influencing antibody recognition for histones because of the presence of extensive modifications on histone tails, therefore future studies may benefit from additional mass spectrometry analyses.(42, 43)

While this study is the first evaluation of H4K20me3 abundance in models of cardiac stress, two previous studies have reported changes in H4K20me3 in the cardiomyocyte during targeted analyses of specific pathways utilizing gain- and loss-of-function. The first study, by Oyama et al., showed that cardiac-specific deletion of the heterochromatin protein 1 gamma (HP1γ), which typically forms heterochromatin by associating with H3K9me3, had no effect on H3K9me3 levels but decreased H4K20me3 abundance in isolated cardiomyocytes.(52) The second study, by Guan et al., investigated the TGF-β signaling pathway in cardiomyocytes and showed that TGF-β signaling leads to miR29-mediated inhibition of the methyltransferase Suv4-20h and resulted in decreased H4K20me3 and decreased cardiac function.(53) Thus, these studies begin to elucidate the possible mechanisms regulating H4K20Me3 in the cardiomyocyte and further link decreased H4K20 trimethylation with cardiac dysfunction.

## AUTHOR CONTRIBUTIONS

S.H, K.D. and S.F. came up with the concept of this manuscript; S.H. and S.F. designed experiments; S.H., M.M., H.S., R.B, S.V. performed experiments and analyzed data; K.D. and S.F. wrote the manuscript. S.H. and K.D. contributed equally to the manuscript. All authors have read and approved the final version of the manuscript.

